# Macaque V1 responses to 2^nd^-order contrast-modulated stimuli and the possible subcortical and cortical contributions

**DOI:** 10.1101/2021.01.28.428627

**Authors:** Nian-Sheng Ju, Shu-Chen Guan, Shi-Ming Tang, Cong Yu

## Abstract

V1 neurons as linear filters supposedly only respond to 1^st^-order luminance-modulated (LM) stimuli, but not 2^nd^-order contrast-modulated (CM) ones. To solve this difficulty, filter-rectify-filter models are proposed, in which first-stage filters respond to CM stimulus elements, and the nonlinear-rectified outputs are summed by a second-stage filter for CM stimulus representation. Correspondingly, neurophysiological evidence shows V1/A17 neurons less responsive to CM stimuli than V2/A18 neurons. Here we used two-photon calcium imaging to demonstrate substantial V1 responses to CM gratings with unimodally distributed LM/CM preferences. Moreover, LM responses are suppressed by LM and CM adaptations regardless of orientation, but CM responses are more suppressed by same-orientation LM and CM adaptations than by orthogonal ones. While LM adaptation results agree with the Hubel-Wiesel view of LGN contributions to V1 orientation responses, CM adaptation results, which include both orientation-unspecific and specific components, may suggest similar subcortical contributions plus additional refinement by recurrent intracortical interactions.

## Introduction

V1 neurons are often modeled as linear filters that are selective to the spatial frequencies (SFs) and orientations of stimuli defined by 1^st^-order statistics, such as luminance (Hubel & Wiesel, 1959, 1962; Carandini, Heeger, & Movshon, 1999). One difficulty the linear filter models face is that they cannot predict perception of contour and boundary stimuli defined by 2^nd^-order statistics, such as contrast, which are abundant in natural scenes. For example, a V1 simple cell as a linear filter is supposedly unresponsive to a 2^nd^-order grating shown in Fig. 1A that changes in contrast, but not luminance.

**Figure 1.**
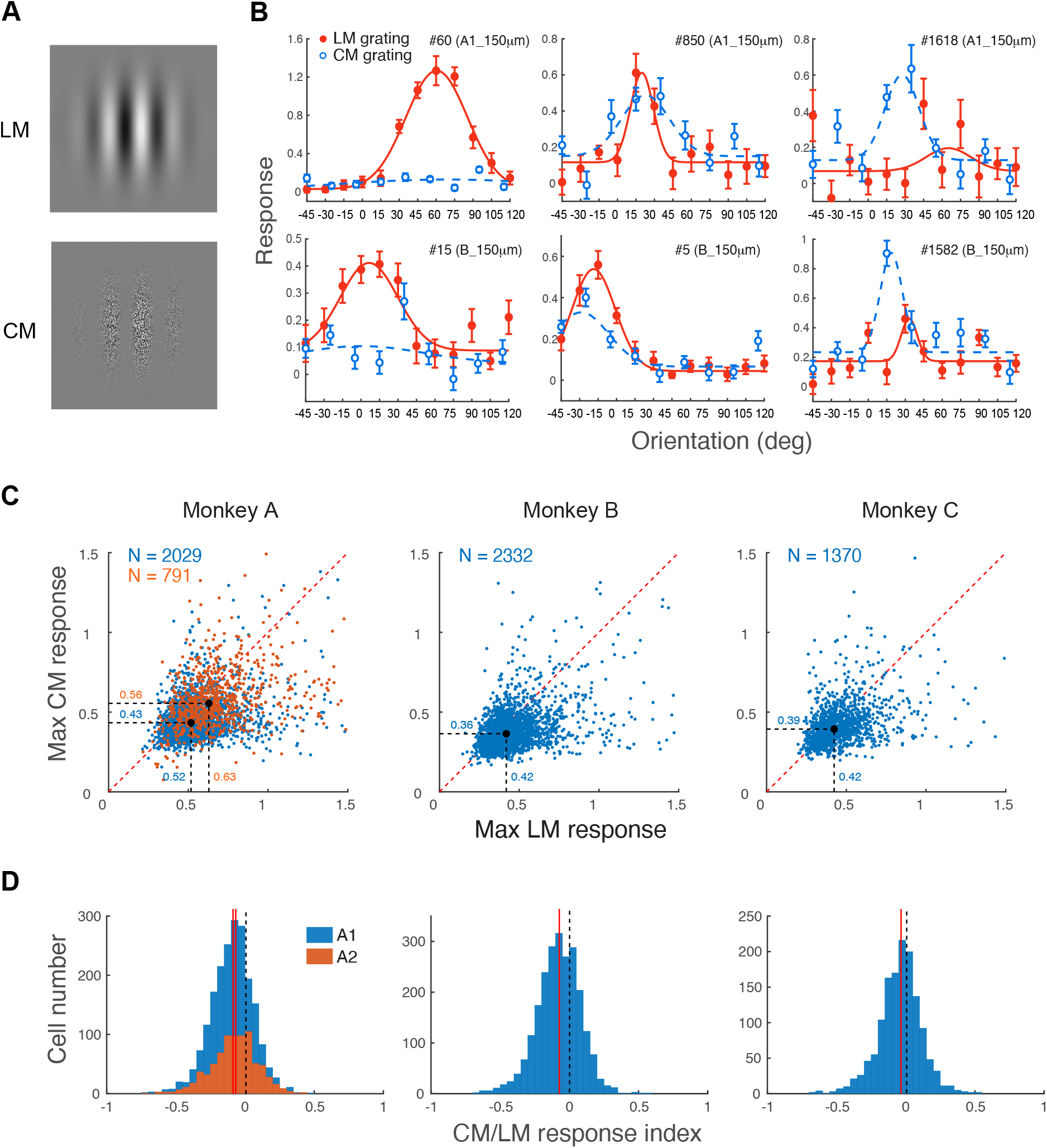
Responses of V1 neurons to LM and CM gratings. **A**. LM (top) and CM (bottom) gratings. The LM grating was a Gabor (a Gaussian-windowed sinusoidal grating), and the CM grating was binary noise multiplied by a Gabor. **B**. Examples of neural responses to LM and CM gratings at various orientations. These neurons showed different preferences to CM and LM gratings. The orientation tuning functions were fitted with a Gaussian function. Error bars indicate ±1 SEM. **C**. Comparisons of maximal CM vs. LM responses of individual neurons in three monkeys. Results at two recording depths of the same recording site in Monkeys A and B were pooled here and later unless otherwise specified. Each dot contrasts one neuron’s CM and LM responses. **D**. Distributions of CM-LM response indices in three monkeys. A neuron would prefer a CM grating more if the index > 0.

To solve this difficulty, various filter-rectify-filter (FRF) models have been proposed (Caelli, 1985; Bergen & Adelson, 1988; Bergen & Landy, 1991; Graham & Sutter, 1998; Landy & Oruc, 2002), in which linear filters first respond to individual stimulus elements (black and white dots in the CM grating of Fig. 1A), then their responses are either half- or full-wave rectified (for example, responses to black dots are either nullified through half-wave rectification or become equivalent to responses to white dots through full-wave rectification, so that responses to black and white dots would not cancel each other), and then the rectified responses are summed by a larger and supposedly lower-SF second-stage linear filter. Often consistent with the FRF models, neurophysiological recording studies have shown that a lot more A18/V2 neurons respond to CM gratings than A17/V1 neurons in cats and monkeys (Zhou & Baker, 1994; El-Shamayleh & Movshon, 2011; G. Li et al., 2014), as if A17/V1 neurons serve as initial linear filters in FRF models and their rectified responses are summed by downstream A18/V2 neurons to signal CM gratings. In addition, there is evidence that V1 neurons may signal CM information through surround suppression (H. Tanaka & Ohzawa, 2009; Hallum & Movshon, 2014). Moreover, even Y cells in LGN can show cortical-cell-like CM responses (i.e., selectivity for carrier SF and orientation) at very high SFs above neurons’ first-order passband as a result of response nonlinearity (Demb, Zaghloul, & Sterling, 2001; Rosenberg, Husson, & Issa, 2010), and Y-cell like neurons in cat A18 may receive direct inputs from LGN Y cells and respond to CM gratings (Gharat & Baker, 2017). Surround suppression and Y-cell response nonlinearity appear to be additional mechanisms to go around the dilemma of linear spatial filters dealing with CM stimuli.

In this study we used two-photon calcium imaging to examine whether macaque V1 neurons can respond to CM gratings directly. The CM stimuli we used were binary noise multiplied by a Gabor function (Fig. 1A). Similar CM stimuli are often used in psychophysical and fMRI studies (Nishida, Ledgeway, & Edwards, 1997; Larsson, Landy, & Heeger, 2006; Ashida, Lingnau, Wall, & Smith, 2007) and an intrinsic-signal optical imaging study (An et al., 2014), but are different from contrast envelopes commonly used in neuronal recording studies made by multiplication of a low-SF grating (as the envelope) and a high-SF grating (as the carrier) of different orientations (Zhou & Baker, 1994; El-Shamayleh & Movshon, 2011; G. Li et al., 2014; Gharat & Baker, 2017). Our main concern with contrast envelope stimuli was that two overlapping gratings at different orientations would elicit cross-orientation suppression, which occurs even when two gratings differ greatly in spatial frequency (Morrone, Burr, & Maffei, 1982; Bonds, 1989; DeAngelis, Robson, Ohzawa, & Freeman, 1992). Our own unpublished observations from a separate two-photon imaging study found significant cross-orientation suppression when the spatial frequencies of two gratings differed by up to 4 octaves.

In addition, we suspected that previous single-unit recording results might have also been affected by sampling issues. These single-unit studies used LM stimuli, such as Gabor gratings, to map the receptive fields, which would exclude neurons mainly responding to CM gratings (Baker, 1999). Moreover, they often under-sampled neurons (e.g., 26 macaque V1 neurons in El-Shamayleh and Movshon (2011) and 42 cat A17 neurons in Zhou and Baker (1994)). If only a small percentage of LM neurons also respond to CM gratings, the detection rate would be very low. These sampling limitations are greatly reduced with two-photon calcium imaging which can simultaneously examine hundreds of neurons’ responses to LM and CM gratings.

## Results

We recorded responses of V1 superficial-layer neurons to drifting LM and CM gratings (Fig. 1A) of various spatial frequencies and orientations in three awake macaque monkeys (The orientation responses of the first two monkeys (A & B) to LM gratings have been reported in a separate paper (Ju, Guan, Tao, Tang, & Yu, 2020). They were re-analyzed here in Figs. 1–3 to compare the CM data). Imaging was performed at two cortical depths (150 and 300 μm from the cortical surface) in Monkeys A and B, and at 150 μm in Monkey C. Monkey A had two recording sites (labeled as A1 & A2), and Monkeys B and C each had one recording site. Imaging processing and data analysis (see Materials and Methods) identified 2820 neurons that were tuned to the orientation of LM and/or CM gratings in Monkey A, 2332 neurons in Monkey B, and 1370 neurons in Monkey C.

As shown in Fig. 1B, some neurons mainly preferred LM gratings (i.e., the left column), some preferred both LM and CM gratings (i.e., the middle column), and some mainly preferred CM gratings (i.e., the right column). When the maximal LM and CM responses of each neuron were contrasted (Fig. 1C), the median maximal LM responses were higher than the median maximal CM response in all three monkeys, by 21% (site 1) and 13% (site 2) in Monkey A, 17% in Monkey B, and 8% in Monkey C, suggesting that more neurons preferred LM gratings to CM gratings (Data at two recording depths of the same recording site in Monkeys A and B were similar, and were thus pooled here and later unless otherwise specified). A CM-LM response index (CLI) was calculated for each neuron: CLI = (R_max_CM_ – R_max_LM_) / (R_max_CM_ + R_max_LM_), so that CLI > 0 would indicate CM preference and CLI < 0 would indicate LM preference. Fig. 1D suggests unimodal distributions of CLI distributions in three monkeys with CLI medians biased toward a preference for LM. Nevertheless, the abundant evidence for CM responses was consistent with our suspicion that previous single-unit recording data were affected by combined effects of cross-orientation suppression and sampling issues.

Based on Gaussian fitting of orientation responses, we divided neurons into three categories: LM neurons which were only tuned to LM grating orientations (only R^2^_LM_ > 0.5), CM neurons which were only tuned to CM grating orientations (only R^2^_CM_ > 0.5), and LM+CM neurons which were tuned to both LM and CM grating orientations (R^2^_LM_ > 0.5 & R^2^_CM_ > 0.5). Fig. 2A indicates that 61.4%, 62.8%, and 53.3% of orientation-tuned neurons were LM neurons, 21.1%, 14.2%, and 12.6% were CM neurons, and 17.6%, 23.0, and 34.1% were LM+CM neurons in Monkeys A, B and C, respectively. Therefore, more than half of the neurons were LM neurons, roughly 20-30% were LM+CM neurons, and 15-20% were CM neurons. The CM neurons would be undetected by conventional RF mapping.

**Figure 2.**
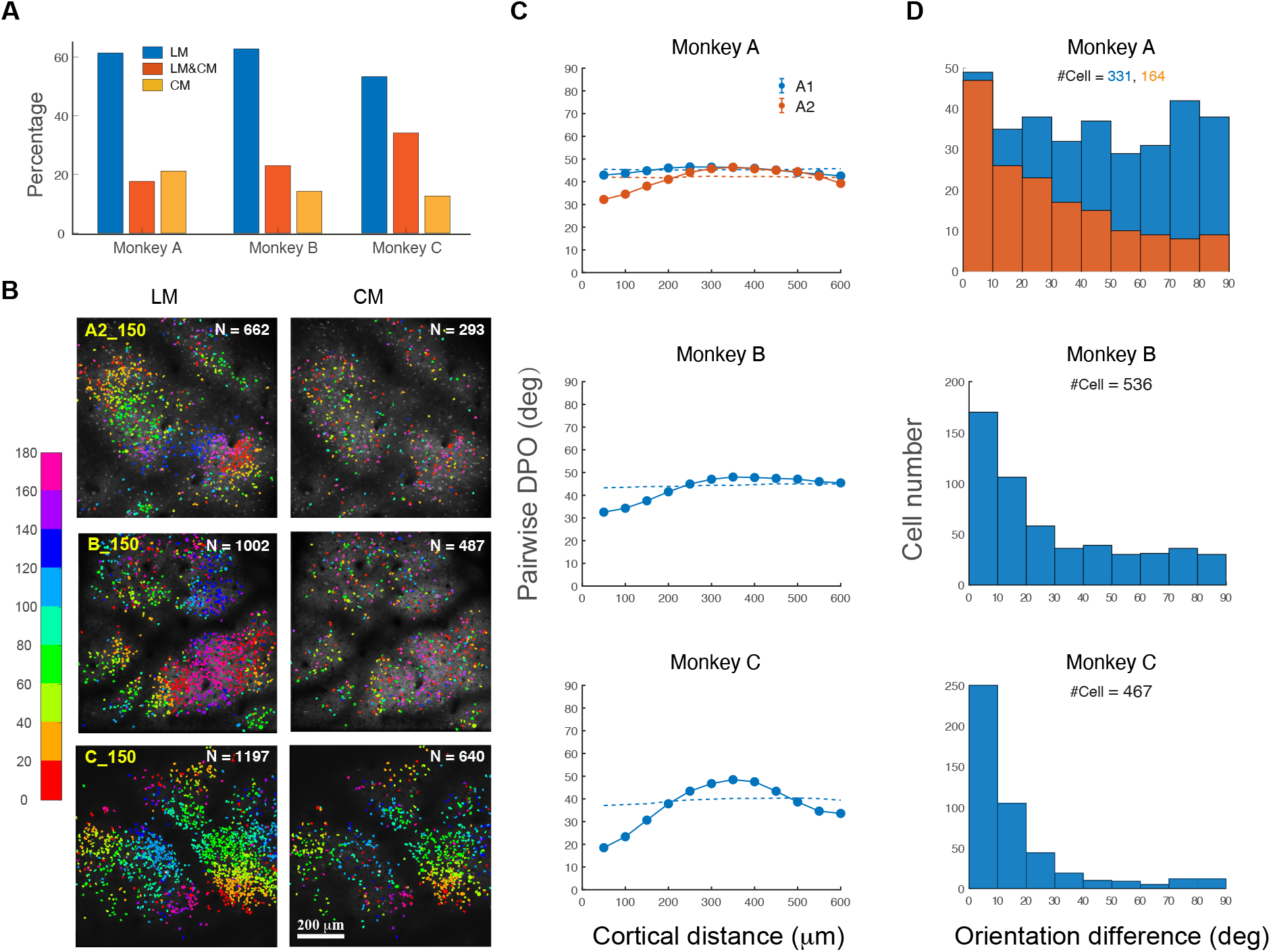
LM and CM orientation tuning. **A**. Percentages of LM, CM, and LM+CM neurons on the basis of Gaussian fitting in three monkeys. **B**. Examples of functional maps of LM and LM+CM orientation tuning with LM gratings (left panels), and of CM and LM+CM orientation tuning with CM gratings (right panels). **C**. Median pairwise difference of preferred orientations (DPO) from neuron pairs, one from the LM orientation map, and the other from the CM orientation map, as a function of the absolute cortical distance. Each datum indicates the median pairwise DPO within a 50 μm bin up to the datum’s corresponding cortical distance on the x-axis. Each corresponding horizontal line represents the baseline simulated with neurons shuffled in position. **C**. The frequency distributions of LM+CM neurons with various differences of optimal LM and CM orientations.

The LM orientation maps including both LM and LM+CM neurons showed clustering of neurons tuned to similar orientations of LM patterns (Figs. 2B left). The orientation clustering of CM and LM+CM neurons in CM maps (Figs. 2B right) was less clear, partly because of fewer number of neurons, except in Map C at 150 μm where orientation clustering was evident. We were particularly interested in whether orientation tuning was spatially clustered across LM and CM maps. To quantify the clustering, we calculated the differences of preferred orientations (DPOs) between neuron pairs, with each pair consisting of one neuron from the LM orientation map and the other neuron from the CM orientation map, as a function of the absolute cortical distance (Fig. 2C). The DPOs were then compared to simulated baselines with all neurons’ positions shuffled in the maps, which should be flat lines around 45° when the orientation preferences of shuffled neurons were evenly distributed (Fig. 2C). The DPOs were similar to baselines in A1, but were lower than the baselines in A2, B and C within a cortical distance of 200 μm. A clustering index (CI) of LM and CM orientation was defined as the inverse of the DPO_mean_/DPO_baseline_ within the first 50 μm of cortical distance. Here the CIs were 1.06 and 1.30 for A1 and A2, 1.33 for B, and 2.00 for C, suggesting little LM and CM orientation clustering in A1, weaker clustering in A2 and B, and stronger clustering in C (Fig. 2C).

Previous single-unit evidence has shown that the same V2 neurons could have very different orientation tuning for CM and LM stimuli (El-Shamayleh & Movshon, 2011). We calculated the differences of preferred LM and CM grating orientations by LM+CM neurons in V1 (Fig. 2D). 36.9% and 58.5% of LM+CM neurons in A1 and A2, 62.3% in B, and 85.7% in C at 150 μm, had their preferred LM and CM orientations differing by 30° or less. Except in A1 where the tuning differences were nearly evenly distributed, in other maps the number of neurons decreased at different rates as the LM vs. CM orientation tuning difference increased. Consistent with these trends, the CIs for LM+CM neurons were 1.00 and 1.62 for A1 and A2, 1.47 for B, and 2.51 for C.

For the above neurons showing LM and/or CM orientation tuning (Fig. 2A), we found that their SF preferences were lower with CM gratings than with LM gratings (Fig. 3; Here a small percentage of neurons not tuned to SF were not included in the analysis). The median preferred SFs were 1.75 cpd, 3.18 cpd, and 4.92 cpd with LM responses, and lower at 1.54, 1.18, and 2.47 cpd with CM responses for Monkeys A to C, respectively (Fig. 3A). For the same LM+CM neurons, their median preferred LM and CM SFs were 1.77 vs. 1.60 cpd in Monkey A, 2.87 vs. 1.25 cpd in Monkey B, and 4.79 vs. 1.46 cpd in Monkey C (Fig. 3B).

**Figure 3.**
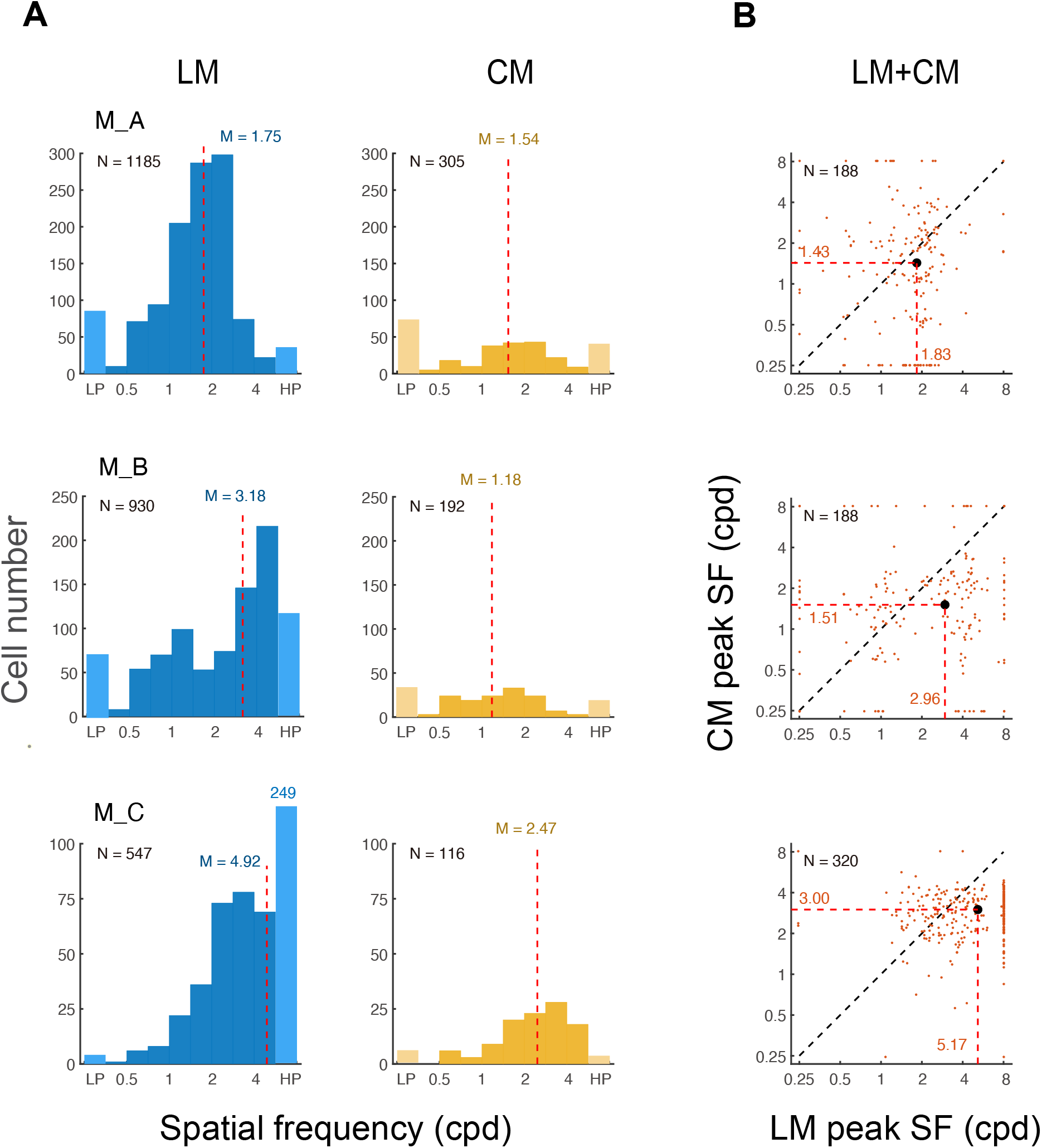
Preferred spatial frequencies of V1 neurons to LM and CM gratings. **A**. Frequency distributions of neurons preferring various spatial frequencies of LM (left) and CM (right) grating. The far left and right bars of lighter bars indicate the number of lowpass and highpass neurons with the current SF measurements (stimulus SFs ranged from 0.25 to 8 cpd), respectively. Vertical dashed lines indicate medians. **B**. Preferred CM spatial frequencies vs. LM spatial frequencies of individual LM+CM neuron. Lowpass and highpass neurons are represented by dots at 0.25 and 8 cpd, respectively. The dashed lines indicate medians.

Psychophysical and fMRI studies have used adaptation experiments to examine the relationship between LM and CM stimulus processing (Nishida et al., 1997; Larsson et al., 2006; Ashida et al., 2007). The general agreement of these studies is that LM and CM stimuli do not interact significantly, and are thus processed in separate channels or pathways. Apparently this finding is contrary to the above two-photon imaging data of a unimodal distribution of LM and CM responses from the same V1 neurons. To explore the relationship between neuronal LM and CM responses, we investigated how adaptation to LM and CM gratings would affect LM and CM responses at the corresponding optimal orientations of neurons. Two conjectures were made. First, if V1 neurons treat LM and CM gratings similarly, as our early data suggested, then the peak response of a specific neuron to one type of stimulus would be at least partially suppressed by adapting to the other type of stimulus at the same orientation. Second, if V1 neuronal responses to CM patterns, like to LM patterns, also have a subcortical origin (Hubel & Wiesel, 1962), or more specifically, if V1 receptive fields take in half-wave rectified responses from aligned LGN neurons to individual black and white dots, then the peak response of a specific neuron to one type of stimulus would be at least partially suppressed by adapting to the other type of stimulus at an orthogonal orientation. This is simply because LGN neurons have little orientation selectivity, and their responses can be suppressed by adaptation at any orientation. This LGN-to-V1 filter-rectify-filter process also applies to the formation of LM orientation according to Hubel and Wiesel (1962) since ON and OFF LGN responses were separately summed.

We recorded the adaptation effects in Monkey C, as well as a new Monkey D. A baseline recording was first performed to measure the neuronal responses to 12 equally spaced orientations at a spatial frequency of 3 cpd on the basis of Monkey C’s earlier comprehensive measurements (3 cpd was the median preferred SF with CM gratings). Then adaptation was measured at 6 of these 12 orientations, again equally spaced. An adaptation stimulus sequence included AAT_n1_AAT_n2_AAT_n3_AAT_n4_ for 20 repeats, where A was the adaptor, and the test stimuli T_n1-n4_ were 4 combinations of LM and CM gratings at the same (0) and orthogonal (90) orientations in a random order. Because each stimulus was presented for 1 s with a 1.5-s inter-stimulus interval, a specific test stimulus was repeated every 30 s, which minimized self-adaptation.

The LM responses were similarly suppressed by both LM and CM adaptors at both the same and orthogonal orientations (Fig. 4A). The adaptation effect was indexed as AEI = (R_peak – R) / (R_peak – R_ortho), where R_peak and R_ortho were pre-adaptation responses to the LM (here) or CM (next paragraph) test stimuli at the preferred and orthogonal orientations, respectively, and R was the post-adaptation response to the LM or CM test stimuli at the preferred orientation. AEI = 0 would indicate no adaptation effect (no reduction of neural response), AEI = 1 would indicate that the neuronal response at the optimal orientation was reduced to the pre-adaptation response level at an orthogonal orientation, and AEI > 1 would indicate that the post-adaptation neuronal response was suppressed to be below the pre-adaptation response level at an orthogonal orientation. Here the median AEIs for LM responses were 0.52, 0.54, 0.67, and 0.77 by adaptors LM/0, CM/0, LM/90, and CM/90, respectively, in Monkey C, and 0.49, 0.40, 0.39, and 0.39 by adaptors LM/0, LM/90, CM/0, and CM/90, respectively, in Monkey D (Fig. 4B). There was no significant difference among different adaptor conditions including LM and CM at the same and orthogonal orientations (F_3, 1389_ = 2.045, p = 0.106, repeated-measures ANOVA)

**Figure 4.**
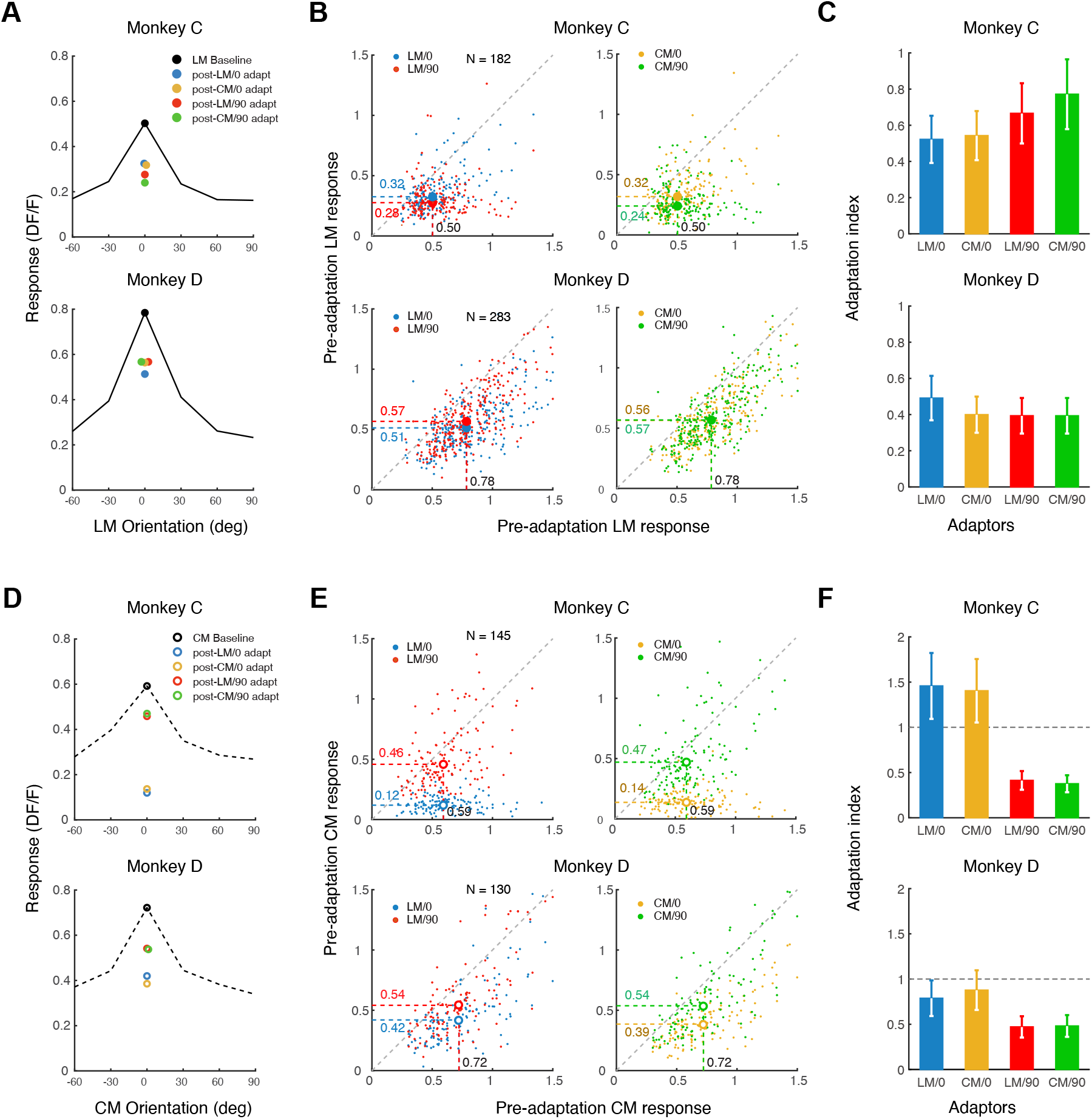
The effects of adaptation on V1 responses to LM and CM gratings. **A**. The median orientation tuning function of neurons with LM gratings and the median responses at the peak orientation before adaptation and after LM/0, CM/0, LM/90, or CM/90 adaptation in two monkeys. Some overlapping data points were slightly laterally shifted for easy viewing. **B**. Scatterplots of individual neuron’s LM responses at the peak orientation before and after adaptation to 4 types of adaptors. The dashed lines indicate median responses. Data points below the diagonal line indicate suppressed responses after adaptation. **C**. The adaptation indices for LM responses with 4 types of adaptors. Error bars show 25 and 75 percentiles. **D**. The median orientation tuning function of neurons with CM gratings and the median responses at the peak orientation before adaptation and after LM/0, CM/0, LM/90, or CM/90 adaptation in two monkeys. **E**. Scatterplots of individual neuron’s CM responses at the peak orientation before and after adaptation to 4 types of adaptors. **F**. The adaptation indices for CM responses with 4 types of adaptors. Error bars show 25 and 75 percentiles.

In contrast, the CM responses were more suppressed by LM and CM adaptors at the same orientation than at the orthogonal orientation (Fig. 4C). The median AEIs for CM responses were 1.46, 1.40, 0.41, and 0.38 by adaptors LM/0, CM/0, LM/90, and CM/90, respectively, in Monkey C, and 0.79, 0.88, 0.47, and 0.48 by adaptors LM/0, CM/0, LM/90, and CM/90, respectively, in Monkey D. There was a significant main effect of adaptors (F_3, 819_ = 102.678, p < 0.001, repeated-measures ANOVA). A contrast analysis indicated that the AEIs with LM/0 and CM/0 adaptors were significantly higher than those with LM/90 and CM/90 adaptors (p < 0.001), while there was no significant difference between LM and CM adaptations at the same orientations (p = 0.637).

One longstanding concern with CM stimuli is that monitor calibration may not get rid of screen luminance nonlinearity completely, which would generate low-contrast luminance cues in CM stimuli, so that the responses of LM neurons to these cues may be mistakenly interpreted as CM responses (Zhou & Baker, 1994). Such a possibility may not fully account for our CM responses because many CM neurons are more responsive to CM stimuli than to LM stimuli, which would have not been predicted if these CM neurons were actually LM neurons. To completely address this concern, we ran a control experiment with Monkey E. We compared LM and CM responses at difference SFs while deliberately making the binary noises visible only at half the SFs. Specifically, the binary noise elements were always 1 x 1 pixel in size. When the stimulus SFs were 0.25, 0.5 and 1 cpd, the viewing distance was 45 cm, so that the pixel size was 2.36 arcmin, which was equal to a maximal SF of 12.7 cpd (the actual SFs in local areas could be lower when more than one pixels of the same polarity were connected). However, for SFs at 2, 4 and 8 cpd, the viewing distance was quadrupled (180 cm), so that the pixel size was 0.59 arcmin, and the equivalent SF was up to 50.8 cpd and invisible at parafovea. What remained with stimuli at these higher SFs would be only potential low-contrast LM cues.

As the SF tuning functions of 4 example neurons (Fig. 5A) suggested, CM stimuli at 28 cpd elicited little neuronal responses. As a result, these neurons preferred the CM stimuli at approximately 1 cpd. In contrast, the same neurons responded strongly to LM stimuli at 2 and 4 cpd, with 2 cpd close to the preferred SF of Neurons #72 and #218. These individual results were confirmed by the difference in frequency distributions of preferred SFs in LM (median SF = 2.35 cpd) and CM neurons (median SF = 0.91 cpd) (Fig. 5B). These results provided direct evidence that our CM data were little affected by screen luminance nonlinearity.

**Figure 5.**
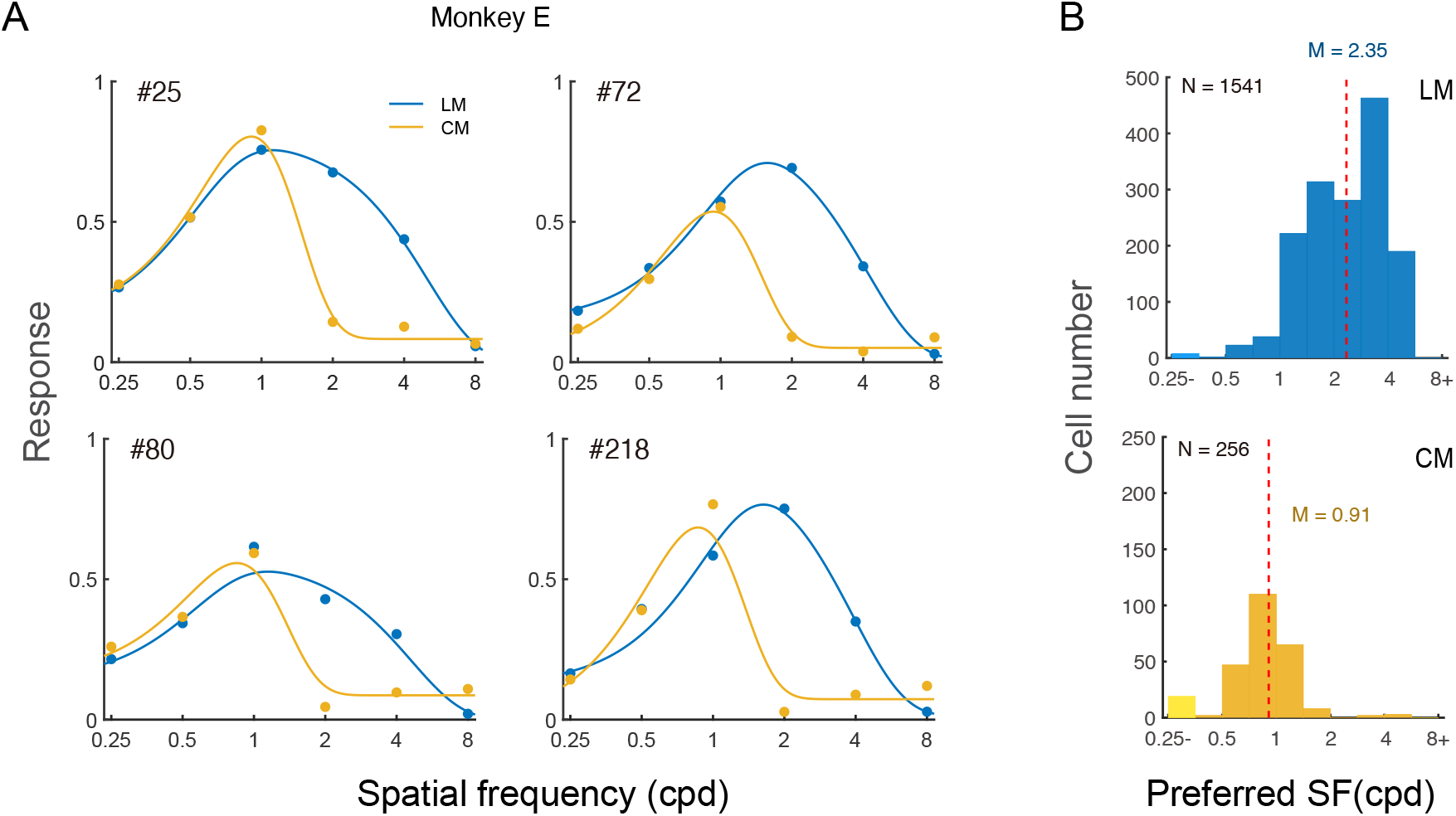
The effects of potential monitor nonlinearity on CM responses. **A**. The SF tuning functions of 4 example neurons with LM and CM stimuli, respectively. The noise elements of CM stimuli were mostly invisible at 2, 4, and 8 cpd. **B**. Frequency distributions of detected LM (left) and CM (right) neurons against preferred SFs. The vertical dashed lines indicate median preferred SFs.

## Discussion

Our results reveal that V1 superficial-layer cells respond to both LM and CM stimuli (Figs. 1B, C), with LM and CM preferences forming a unimodal distribution that skews towards LM stimuli (Fig. 1D). This finding is in agreement with an intrinsic-signal optical imaging study showing V1 responses to both LM and CM stimuli (An et al., 2014). More than half of the neurons are tuned to only LM-orientations, less (20%-50%) are to both LM and CM orientations, and the least (<20%) to only CM orientations (Fig. 2A). The median preferred SFs with CM stimuli are lower than those with LM stimuli and are within the first-order SF passband (Fig. 3), indicating that these CM responses may not reflect response nonlinearity that leads to CM responses at high SFs beyond neurons’ first-order passbands (Gharat & Baker, 2017). As pointed out earlier, our stimulus design prevents the explanation that these CM responses reflect surround suppression associated with oriented carrier grating in contrast envelope stimuli (H. Tanaka & Ohzawa, 2009; Hallum & Movshon, 2014).

Adaptation evidence may shed lights on the mechanisms underlying neuronal LM and CM responses. As Figs. 4A-C shows, neurons’ peak LM responses are similarly suppressed by LM and CM adaptors at both the same and orthogonal orientations, which is reminiscent of the classical view that V1 simple cell receptive fields pool outputs from aligned ON- or OFF-center LGN cell receptive fields (Hubel & Wiesel, 1962; K. Tanaka, 1985; Reid & Alonso, 1995). Here simple cells’ pooling of the same-sign LGN inputs suggests FRF processing in that simple cells sum half-wave rectified outputs of LGN cells. Because LGN neurons have little orientation preferences, LM adaptors and the bright and dark elements of CM adaptors regardless of orientation can suppress the responses of aligned ON- and OFF-center LGN neurons (Shou, Li, Zhou, & Hu, 1996) that continue to form simple cell receptive fields in V1. Additional roles of recurrent intracortical excitation and inhibition that has been proposed to sharpen orientation tuning (Ben-Yishai, Bar-Or, & Sompolinsky, 1995; Douglas, Koch, Mahowald, Martin, & Suarez, 1995; Somers, Nelson, & Sur, 1995) appear to be minimal with the current high contrast stimuli because of similar adaptation effects at the same and orthogonal orientations.

CM responses may be similarly explained with additionally more intracortical contributions. V1 simple cells may first sum half-wave rectified inputs from aligned ON- or OFF-center LGN neurons that are activated by CM stimuli. However, the resulting orientation tuning of simple cells is likely to be poorer because of insufficient summation of LGN inputs activated by CM gratings than by LM gratings (the perceived contrast of a CM grating is also lower than that of a LM grating with equal physical contrast (Fig. 1A)). It is under this circumstance that the proposed intracortical interactions, especially orientation-specific intracortical excitations (Ben-Yishai et al., 1995; Douglas et al., 1995; Somers et al., 1995) may play an important role by further sharpening CM orientation tuning. Accordingly, adaptation at optimal orientations suppresses orientation-unspecific subcortical contributions to CM responses (indicated by the suppression indices with LM/90 and CM/90 conditions in Fig. 4F), as well as orientation specific intracortical contributions (The suppression indices of LM/0 and CM/0 subtracted by indices of LM/90 and CM/90 in Fig. 4F). This double suppression may lead to higher CM response suppression than LM response suppression, although we do not have an explanation why in Monkey C the CM responses were suppressed to be below the pre-adaptation orthogonal responses (Figs. 4C vs. 4F). Apparently this ad hoc analysis does not introduce special mechanisms for V1 CM processing, but only involves known LGN response pooling and intracortical interactions, which is similar to the early Bergen and Adelson model on early vision and texture perception (Bergen & Adelson, 1988).

The observation of substantial CM responses of V1 neurons is consistent with our suspicion that the lack of V1 evidence for contrast envelope processing in single-unit studies (Zhou & Baker, 1994; El-Shamayleh & Movshon, 2011) might be related to cross-orientation inhibition. When two gratings at different orientations multiply to form a plaid, cross-orientation inhibition takes effect even when the spatial frequencies of two gratings differ (Morrone et al., 1982; Bonds, 1989; DeAngelis et al., 1992), which may prevent V1 neurons from responding to contrast envelopes. Some V1 neurons actually prefer plaid stimuli to grating stimuli as our recent two-photon imaging evidence suggests (Guan, Zhang, Zhang, Tang, & Yu, 2020). However, these plaid neurons are likely excluded during initial receptive field mapping in neuronal recording studies because plaid neurons tend not to respond much to gratings. One missing link is whether V2 or Area 18, where more CM neurons have been discovered with contrast envelope stimuli (Zhou & Baker, 1994; El-Shamayleh & Movshon, 2011; G. Li et al., 2014), contain substantially more neurons that are responsive to both gratings and plaids. One related concern is whether the CM responses in our study were also affected by cross-orientation inhibition. This is because cross-orientation inhibition may be a result of general contrast gain control, which normalizes the neuron’s tuned response by the sum of all the contrast energy that falls within the classical receptive field (Heeger, 1992). However, such a possibility is not supported by our data. As Fig. 1D shows, the CM/LM response index was larger than −0.1, suggesting mildly weaker CM responses than LM responses. In contrast, the cross orientation inhibition index (calculated in the same manner as the CM/CM response index) was −0.32 with plaid stimuli (multiplication of two orthogonal gratings) in Guan et al. (2020), which suggests significantly stronger cross orientation inhibition. Therefore, V1 responses to CM stimuli with noise carriers were much less affected by cross-orientation inhibition than responses to plaid stimuli.

Finally, we want to point out that our analysis cannot explain why neuronal adaptation evidence, which suggests a more upstream FRF processing from LGN to V1 (see earlier discussion), is different from psychophysical (Nishida et al., 1997) and fMRI adaptation evidence (Nishida et al., 1997; Larsson et al., 2006; Ashida et al., 2007). Assuming these early studies had not been affected by the small number of participants (e.g., total N = 14 in three cited fMRI studies), their conclusion is at odds with the FRF models in general by suggesting independent LM and CM processing. As some fMRI evidence indicates, CM processing involved higher brain areas such as V3, V4, and MT+ (Larsson et al., 2006; Ashida et al., 2007). One possibility is that psychophysical and fMRI adaptation experiments actually have targeted later stages of CM processing. For example, high brain areas may process LM and CM stimuli independently on the basis of their appearance differences.

## Materials and Methods

### Monkey preparation

Monkey preparations were identical to what reported in a previous paper (Guan et al., 2020). Five macaque monkeys (aged 5-8 years) were each prepared with two sequential surgeries under general anesthesia and strictly sterile conditions. In the first surgery, a 20mm diameter craniotomy was performed on the skull over V1. The dura was opened and multiple tracks of 100-150 nL AAV1.hSynap.GCaMP5G.WPRE.SV40 (AV-1-PV2478, titer 2.37e13 (GC/ml), Penn Vector Core) were pressure-injected at a depth of ~350 μm. The dura was then sutured, then the skull cap was re-attached with three titanium lugs and six screws, and the scalp was sewn up. After the surgery, the animal was returned to the cage, treated with injectable antibiotics (Ceftriaxone sodium, Youcare Pharmaceutical Group, China) for one week. The second surgery was performed 45 days later. A T-shaped steel frame was installed for head stabilization, and an optical window was inserted onto the cortical surface. More details of the preparation and surgical procedures can be found in M. Li, Liu, Jiang, Lee, and Tang (2017). The procedures were approved by the Institutional Animal Care and Use Committee, Peking University.

### Behavioral task

After a ten-day recovery from the second surgery, monkeys were seated in primate chairs with head restraint. They were trained to hold fixation on a small white spot (0.1°) with eye positions monitored by an ISCAN ETL-200 infrared eye-tracking system (ISCAN Inc.) at a 120-Hz sampling rate. During the experiment, trials with the eye position deviated 1.5° or more from the fixation were discarded as ones with saccades and repeated. For the remaining trials, the eye positions were mostly concentrated around the fixation point, with eye positions in over 95% of trials within 0.5° from the fixation point. The viewing was binocular.

### Visual stimuli

For Monkeys A, B, D, and E, visual stimuli were generated by the ViSaGe system (Cambridge Research Systems) and presented on a 21’’ Sony G520 CRT monitor with a refresh rate of 80 Hz. Monitor resolution was set at 1280 pixel × 960 pixel. The pixel size was 0.31 mm × 0.31 mm. Because of the space limit, viewing distances varied depending on the stimulus spatial frequency (LM gratings: 30 cm at 0.25/0.5/1 cpd, 60 cm at 2cpd, and 120 cm at 4/8 cpd; CM gratings: 30 cm at 0.25/0.5/1 cpd, 60 cm at 2/4 cpd, and 120 cm at 8 cpd), except for Monkey E the view distances were 0.45 cm at 0.25/0.5/1 cpd and 1.80 cm at 2/4/8 cpd (see Fig. 5). For Monkey C, visual stimuli were generated by Psychotoolbox 3 (Pelli & Zhang, 1991) and presented on a 27’’ Acer XB271HU LCD monitor. The refresh rate of the monitor was native at 80 Hz and the resolution was native at 2560 pixel × 1440 pixel. The pixel size was 0.23 mm × 0.23 mm. The viewing distance was 50 cm for lower frequencies (0.25 - 2 cpd) and 100 cm for higher frequencies (4 and 8 cpd). For both monitors, the screen luminance of two monitors was linearized by an 8-bit look-up table, and the mean luminance was ~47 cd/m2.

A drifting square-wave grating (3 cycles/s, full contrast, 4 cpd spatial frequency, and 0.4° diameter in size) was first used to determine the population receptive field size and location associated with a recording site (2-4° eccentricity), as well as ocular dominance columns when monocularly presented to confirm the V1 location. This process was quick with the use of a 4 × objective lens mounted on the two-photon microscope, which revealed no cell-specific information.

Cell-specific responses were then measured with a high-contrast (0.9) LM stimulus, which was a Gabor gating (Gaussian-windowed sinusoidal grating) drifting at 2 cycles/s in opposite directions perpendicular to the Gabor orientation, or a CM stimulus, which was the Gabor grating multiplied by binary noise of 1 or −1. The Gabor grating varied at 12 equal-spaced orientations from 0° to 165° in 15° steps, and 6 spatial frequencies from 0.25 to 8 cpd in 1 octave steps. In addition, our pilot measurements suggested very strong surround suppression with larger stimuli. Therefore, we used three stimulus sizes at each spatial frequency, so as to estimate the best responses of each neuron with the most center summation and least surround suppression. Specifically, the σ of the Gaussian envelope of the Gabor were 0.64λ and 0.85λ at all SFs, and was additionally smaller at 0.42λ when SFs were 0.25, 0.5, and 1 cpd, and larger at 1.06λ when SFs were 2, 4, and 8 cpd (λ: wavelength; Gabors with the same σ in wavelength unit had the same number of cycles). Here at the smallest σ (0.42λ), the Gabors still had sufficient number of cycles (frequency bandwidths = 1 octave) (Graham, 1989), so that the actual stimulus SFs were precise at nominal values. For CM Gabors, to ensure visibility of noise elements at all spatial frequencies, the noise element size was 1×1 pixel at 0.25/0.5/1 cpd, 2×2 pixel at 2/4 cpd, and 3×3 pixel at 8 cpd when the CRT monitor was used, and 1×1 pixel at 0.25/0.5/1 cpd and 2×2 pixel at 2/4/8 cpd when the LCD monitor was used.

The stimuli at a specific viewing distance were pseudo-randomly presented. Each stimulus was presented for 1 s, with an inter-stimulus interval (ISI) of 1500 ms that is sufficient to allow the calcium signals back to the baseline level (Guan et al., 2020). Each stimulus condition was repeated 12 times with half trials for each opposite direction. Imaging of all orientations, spatial frequencies, and stimulus sizes for either LM or CM stimuli at a specific recording site and depth was completed in one session that lasted 3 - 4 hours.

### Two-photon imaging

Two-photon imaging was performed with a Prairie Ultima IV (In Vivo) two-photon microscope (Prairie Technologies) (Monkeys A, B, D) or a FENTOSmart two-photon microscope (Femtonics) (Monkey C), and a Ti:sapphire laser (Mai Tai eHP, Spectra Physics). One or two windows of 850 x 850 μm^2^ were selected in each animal and imaged using 1000-nm femtosecond laser under a 16 × objective lens (0.8 N.A., Nikon) at a resolution of 1.6 μm/pixel. Fast resonant scanning mode (32 fps) was chosen to obtain continuous images of neuronal activity (8 fps after averaging every 4 frames). For either LM or CM stimuli, recordings at two depths of the same site were completed in two consecutive days. On the first day, recordings were performed at 150 μm with LM stimuli. Some neurons with high brightness or unique dendrite patterns were selected as landmarks. In later recording sessions, the same field of view (FOV) at 150 μm was first located with the help of landmark neurons. Then the depth plane was lowered to 300 μm when necessary. LM and CM recordings were separated by 1-5 days.

### Imaging data analysis: Initial screening of ROIs

Data were analyzed with customized MATLAB codes. A normalized cross-correlation based translation algorithm was used to reduce motion artifacts (M. Li et al., 2017). Then fluorescence changes were associated with corresponding visual stimuli through the time sequence information recorded by Neural Signal Processor (Cerebus system, Blackrock Microsystem). By subtracting the mean of the 4 frames before stimuli onset (*F0*) from the average of the 6th-9th frames after stimuli onset (*F*) across 5 or 6 repeated trials for the same stimulus condition (same orientation, spatial frequency, size, and drifting direction), the differential image (*ΔF = F - F0*) was obtained.

The regions of interest (ROIs) or possible neurons were decided through sequential analysis of 512 differential images in the order of stimuli type (2), SF (6), size (3), and orientation (12) (2 x 6 x 3 x 12 = 512). The first differential image was filtered with a bandpass Gaussian filter (size = 2-10 pixels), and connected subsets of pixels (>25 pixels, which would exclude smaller vertical neuropils) with average pixel value > 3 standard deviations of the mean brightness were selected as ROIs. Then the areas of these ROIs were set to mean brightness in the next differential image before the bandpass filtering and thresholding were performed (This measure would eventually reduce the SDs of differential images and facilitate detection of neurons with relatively low fluorescence responses). If a new ROI and an existing ROI from the previous differential image overlapped, the new ROI would be on its own if the overlapping area OA < 1/4 ROI_new_, discarded if 1/4 ROI_new_ < OA < 3/4 ROI_new_, and merged with the existing ROI if OA > 3/4 ROI_new_. The merges would help smooth the contours of the final ROIs. This process went on through all 512 differential images twice to select ROIs. Finally, the roundness for each ROI was calculated as:

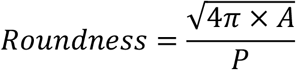

where *A* was the ROI’s area, and *P* was the perimeter. Only ROIs with roundness larger than 0.9, which would exclude horizontal neuropils, were selected for further analysis.

### Imaging data analysis: Orientation-/SF-tuned neurons

The ratio of fluorescence change (*ΔF/F0*) was calculated as a neuron’s response to a specific stimulus condition. For a specific cell’s response to a specific stimulus condition, the *F0_n_* of the n-th trial was the average of 4 frames before stimulus onset, and F_n_ was the average of 5th-8th or 6th-9th frames after stimulus onset, whichever was greater. F0_n_ was then averaged across 10 or 12 trials to obtain the baseline F0 for all trials (for the purpose of reducing noises in the calculations of responses), and ΔF_n_/F_0_ = (F_n_-F0)/F0 was taken as the neuron’s response to this stimulus at this trial. For a small portion of neurons (e.g., ~3% in Monkeys A and B when responding to LM Gabors) showing direction selectivity as their responses to two opposite directions differed significantly (p < 0.05, Friedman test), the 5-6 trials at the preferred direction was considered for calculations of △F_n_/F0 as the cell’s responses to a particular stimulus. F0 was still averaged over 10-12 trials at two opposite directions.

Several steps were then taken to decide whether a neuron was tuned to orientation and/or SF of LM or CM stimuli. First, the spatial frequency, orientation, and size producing the maximal response among all LM or CM conditions were selected. Then responses to other 11 orientations were decided at the selected spatial frequency and size. Second, to select orientation tuned neurons, a non-parametric Friedman test was performed to test whether a neuron’s responses at 12 orientations were significantly different from each other. To reduce Type-I errors, the significance level was set at α = 0.01. Third, for those showing significant difference, the orientation tuning function of each neuron was fitted with a Gaussian model:

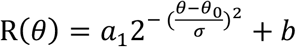

where R(θ) was the response at orientation θ, free parameters a_1_, θ_0_, σ, and b were the amplitude, peak orientation, standard deviation of the Gaussian function, and minimal response of the neuron, respectively. Only neurons with goodness of fit R^2^ > 0.5 were finally selected as orientation tuned neurons (Fig. 1B). The amplitude parameter a1 was positive in all selected orientation neurons. Fourth, the SF tuning function of each neuron was fitted with a Different-of-Gaussian model.

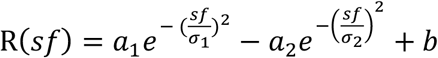

where R(sf) was a neuron’s response at spatial frequency sf, free parameters a_1_, σ_1_, a_2_, and σ_2_ were amplitudes and standard deviations of two Gaussians, respectively, and b was the minimal response among 6 spatial frequencies. Only neurons with goodness of fit R^2^ > 0.5 were finally selected as SF tuned neurons.

## Acknowledgments

This study was supported by Natural Science Foundation of China grants 31230030 and 31730109, and funds from Peking-Tsinghua Center for Life Sciences, Peking University.

